# Genetic and transcriptomic dissection of nitrate-independent function of Arabidopsis NRT1.1/NPF6.3/CHL1 under high ammonium condition

**DOI:** 10.1101/2023.10.17.562629

**Authors:** Takushi Hachiya, Nobue Makita, Liên Bach, Alain Gojon, Tsuyoshi Nakagawa, Hitoshi Sakakibara

## Abstract

The Arabidopsis nitrate transceptor NRT1.1/NPF6.3/CHL1 regulates physiological responses to nitrate. Several studies have reported that Arabidopsis plants lacking NRT1.1 show enhanced shoot growth under toxic levels of ammonium without nitrate, suggesting a nitrate-independent function for NRT1.1. To further investigate this nitrate-independent function and its impact on ammonium tolerance, we conducted genetic analysis, tissue-specific expression analysis, and transcriptome analysis using various NRT1.1-related lines. Transgenic plants expressing either nonphosphomimic or phosphomimic mutants of NRT1.1 exhibited similar ammonium tolerance to the wild-type. The *chl1-9* mutant, in which NRT1.1 with the P492L substitution is localized intracellularly rather than at the plasma membrane and fails to transport nitrate, showed significantly improved ammonium tolerance. Confocal imaging revealed that the NRT1.1-GFP signal was detected in the plasma membrane of various tissues, including cotyledon pavement cells, hypocotyl epidermal cells, mesophyll cells, root cap cells, and epidermal cells near root tips. In early seedlings, the absence of functional NRT1.1 altered the expression of genes associated with aliphatic glucosinolate biosynthesis, ethylene signaling, and low pH stress. Genes predicted to encode products localized to the extracellular space were enriched among those differentially expressed due to *NRT1.1* deficiency. Our data suggest that in the absence of nitrate, plasma membrane-targeted NRT1.1 reduces ammonium tolerance irrespective of its phosphorylation state with alterations of gene expression associated with stress and senescence.

## 1. Introduction

Most land plants primarily rely on soil nitrate as their main nitrogen (N) source. Nitrate serves not just as a substrate for N assimilation but also as a vital signaling molecule, regulating genes related to nitrogen acquisition and root system development (Wang et al. 2004; Okamoto et al. 2019). These nitrate responses require molecular components to perceive nitrate and to drive nitrate-dependent signaling. The Arabidopsis nitrate transporter NRT1.1/NPF6.3/CHL1 plays a crucial role as a nitrate sensor, orchestrating various plant responses to nitrate supply (Remans et al. 2006; Ho et al. 2009; Bouguyon et al. 2015). Downstream of NRT1.1, calcium ions serve as secondary messengers, stimulating the expression of nitrate-responsive genes, including the major high-affinity nitrate transport gene *NRT2.1* (Riveras et al. 2015). Recent findings indicate that the subgroup III protein kinase CPK perceives calcium signals and phosphorylates the transcription factor NLP7 (Liu et al. 2017), which directly binds intracellular nitrate, regulating nitrate-dependent gene expression (Liu et al. 2022). Furthermore, NRT1.1 enhances the expression of the transcription factor *ANR1* and the *AFB3* auxin receptor gene in roots, promoting lateral root development and elongation in response to nitrate supply (Remans et al. 2006; Vidal et al. 2010). The magnitude of these NRT1.1-dependent responses is finely tuned by the phosphorylation status of NRT1.1 at the T101 residue (Ho et al. 2009; Bouguyon et al. 2015).

In addition to its nitrate-dependent functions, NRT1.1 exhibits significant roles in the absence of nitrate. It aids the basipetal transport of auxin out of the lateral root primordia when nitrate is absent (Krouk et al. 2010). Exogenous nitrate inhibits auxin transport, leading to auxin accumulation in primordia and the stimulation of lateral root emergence in nitrate-rich patches (Mounier et al. 2014). Walch-Liu and Forde (2008) observed that exogenous glutamate inhibits primary root elongation at the root tip, a process antagonized by nitrate presence, depending on phosphorylated NRT1.1. Intriguingly, overexpression of nonphosphomimetic NRT1.1 (T101A) heightens glutamate sensitivity in primary roots even without nitrate. Furthermore, studies including ours have demonstrated that the absence of functional NRT1.1 significantly alleviates growth suppression and chlorosis under toxic ammonium levels as the sole N source (Hachiya et al. 2011; Jian et al. 2018; Liu et al. 2020). NRT1.1 promotes ammonium accumulation, altering ammonium metabolism and subsequently upregulating ethylene signaling, resulting in ammonium toxicity (Jian et al. 2018). Moreover, under high ammonium and NaCl concentrations without nitrate, NRT1.1 transports chloride, leading to chloride accumulation, especially in roots (Liu et al. 2020). This chloride accumulation contributes to ammonium toxicity under high salt conditions.

The aforementioned observations clearly indicate a nitrate-independent function for NRT1.1, although the precise mechanism remains elusive. This study aimed to comprehensively uncover the primary nitrate-independent function of NRT1.1 in ammonium tolerance. Employing various NRT1.1-related lines, we conducted genetic analyses, tissue-specific expression studies using quantitative PCR (qPCR) and confocal microscopy, and transcriptome analyses. Our findings reveal that (i) plasma membrane-targeted NRT1.1 diminishes ammonium tolerance regardless of its phosphorylation state, (ii) NRT1.1 is expressed across most seedling tissues, and (iii) NRT1.1 primarily modulates the expression of genes associated with aliphatic glucosinolate (GSL) biosynthesis, ethylene signaling, and low pH stress.

## 2. Materials and Methods

### 2.1. Plant materials and growth conditions

In this study, Arabidopsis plants were used. The T-DNA insertion mutants *nrt1.1* (Hachiya et al. 2011), *cipk23-4* (Ho et al. 2009), and the gamma ray-mutagenized mutant c*hl1-5* (Tsay et al. 1993a) in the Col background were obtained from the European Arabidopsis Stock Center (NASC). The gamma-ray-mutagenized mutant *chl1-6* in the L*er* background (Tsay et al. 1993b) was sourced from the Arabidopsis Biological Resource Center. The transposon tag line *pst16286* in the Nossen background (Ito et al. 2002; Kuromori et al. 2004) was purchased from the RIKEN Bioresource Center (BRC). The lines harboring the estradiol-inducible *NRT1.1-mCherry* and the *pNRT1.1:NRT1.1-GFPloop* in the *chl1-5* background were used in previous studies (Bouguyon et al. 2015; Bouguyon et al. 2016). The *chl1-9* in the Col background and the *T101D* and *T101A* lines in the *chl1-5* background (Ho et al. 2009) were provided by Dr. Yi-Fang Tsay (Academia Sinica).

The seeds underwent surface sterilization and were placed in plastic Petri dishes (diameter: 90 mm; depth: 20 mm; Iwaki, Tokyo, Japan) with approximately 30 mL of N-modified Murashige and Skoog medium. The medium included 4.7 mM MES-KOH (pH 5.7), 2% (w/v) sucrose, and 0.25% (w/v) gellan gum (Wako, Osaka, Japan). Two different N and K sources were used: 10 mM KNO_3_ (10 mM NO_3_^-^ condition) or 5 mM (NH_4_)_2_SO_4_ with 10 mM KCl (10 mM NH_4_^+^ condition). After being kept in the dark at 4 °C for 3 d, the plants were grown horizontally under a photosynthetic photon flux density of 100–130 μmol m^−2^ s^−1^ (16 h light/8 h dark cycle) at 23 °C. For transfer experiments, surface-sterilized seeds were sown in larger plastic Petri dishes (length: 140 mm; width: 100 mm; depth: 20 mm; Eiken Chemical Co. Ltd., Taito-ku, Tokyo, Japan) containing 50 mL of half-strength modified Murashige and Skoog medium with 2.5 mM ammonium as the sole N source at pH 6.7 (Okamoto et al. 2019). After 3 days in the dark at 4 °C, the plants were grown vertically for 5 d under a photosynthetic photon flux density of 100–130 μmol m^−2^ s^−1^ (16 h light/8 h dark cycle) at 23 °C. The plants were then transferred to different N conditions for subsequent experiments. Further details regarding plantlet cultivation are provided in the Results section and figure legends.

### 2.2. Extraction of RNA

The whole seedlings, shoots, and roots were harvested and promptly frozen in liquid N_2_, then stored at −80 °C until needed. The frozen samples were ground using TissueLyser II (QIAGEN) with zirconia beads (5 mm diameter). Total RNA was extracted using the RNeasy Plant Mini Kit (Qiagen) and treated with on-column DNase digestion following the manufacturer’s instructions.

### 2.3. RT-qPCR

Reverse transcription (RT) was conducted using ReverTra Ace qPCR RT Master Mix with gDNA Remover (Toyobo Co. Ltd., Tokyo, Japan) following the manufacturer’s guidelines. The resulting cDNA was diluted tenfold with distilled water for quantitative PCR (qPCR). Transcript levels were assessed using a StepOnePlus Real-Time PCR System (Thermo Fisher Scientific, Waltham, MA, USA). In the presence of 10-µL KAPA SYBR FAST qPCR Kit (Nippon Genetics Co. Ltd., Tokyo, Japan), 0.4 µL specific primers (0.2 µM final concentration), and 7.2 µL sterile water, 2 µL of obtained cDNA was amplified. *ACTIN3* (Hachiya et al. 2021) served as internal standards. Standard curves were generated using plasmid DNA containing target cDNAs or total cDNAs. Refer to Supplementary Table 1 for primer sequences.

### 2.4. Microarray analysis

Microarray analysis was conducted using 3-day-old (72 h) and 5-day-old (120 h) whole seedlings of Col, *chl1-5*, and *nrt1.1* following the protocol outlined in Hachiya et al. (2021). RNA quality was evaluated using an Agilent 2100 bioanalyzer (Agilent Technologies). RNA amplification, labeling, hybridization, and scanning were performed using the 3′ IVT Express Kit (Affymetrix) and GeneChip Arabidopsis Genome ATH1 Array (Affymetrix) as per the manufacturer’s instructions. Data from the microarray chips were normalized using the Microarray Suite 5.0 (MAS5) method (Affymetrix). Transcripts labeled as “absent” or “marginal” were excluded from subsequent quantitative analysis. The raw microarray data used in this study are accessible in the ArrayExpress database at EMBL-EBI under accession number E-MTAB-13395.

### 2.5. Observation of cotyledon

Bright-field imaging of Arabidopsis early seedling cotyledons was conducted using an all-in-one fluorescence microscope (BZ-X710, KEYENCE, Japan) equipped with a Nikon 40× objective (CFI Plan Apo λ 40×/0.95). To create a high-resolution and wide-area continuous image of the cotyledon, individual images were seamlessly merged using the image joint function of the BZ analyzer software (KEYENCE).

### 2.6. Observation of fluorescent signal

Confocal imaging of NRT1.1-GFPloop and propidium iodide (PI) was performed using a Leica SP5 Confocal Microscope (Leica Microsystems). Roots were stained with a 10 µg mL^−1^ PI solution to outline cell shapes. Mesophyll protoplasts were prepared from cotyledons of 12-day-old plants (Plants grown under 10 mM nitrate for 9 days were transferred to 10 mM ammonium and grown for 3 days.) following the method of Endo et al. (2016). NRT1.1-GFPLoop and PI were excited with 488 nm and 543 nm lasers, and emissions were detected from 500 to 530 nm and 590 to 660 nm, respectively, using a Leica 25× objective (HCX IRAPO L 25×/0.95 WATER).

### 2.7. Statistical analysis

The unpaired two-tailed Welch’s *t*-test and the Tukey–Kramer multiple comparison test were conducted using R software v.2.15.3. Additionally, the two-tailed ratio paired t-test was performed using GraphPad Prism software v.9.3.1, assuming a Gaussian distribution.

## 3. Results

### 3.1. NRT1.1 targeted to the plasma membrane reduces ammonium tolerance almost regardless of its phosphorylation status

In our previous study, *NRT1.1*-deficient mutants in the Col background exhibited enhanced shoot growth under 10 mM ammonium conditions (Hachiya et al. 2011). In this study, we consistently observed larger shoot fresh weights (FWs) in *NRT1.1*-deficient mutants from three Arabidopsis accessions (Col; *chl1-5*, *nrt1.1*, *Ler*; *chl1-6*, and Nossen; *pst16286*) compared to wild-type plants under 10 mM ammonium. Conversely, under 10 mM nitrate conditions, the FWs were lower in these mutants (Figure 1a, b). Intriguingly, the induction of NRT1.1-mCherry in the *chl1-5* background, controlled by β-estradiol (Bouguyon et al. 2015), significantly reduced shoot FW under ammonium, whereas β-estradiol-treated Col plants did not show a change in FW (Figure 1c). Under nitrate, shoot growth was restored by the induction of NRT1.1-mCherry, confirming its functionality (Figure 1d). These findings underscore the significant role of Arabidopsis NRT1.1 in ammonium tolerance.

**Figure 1.**
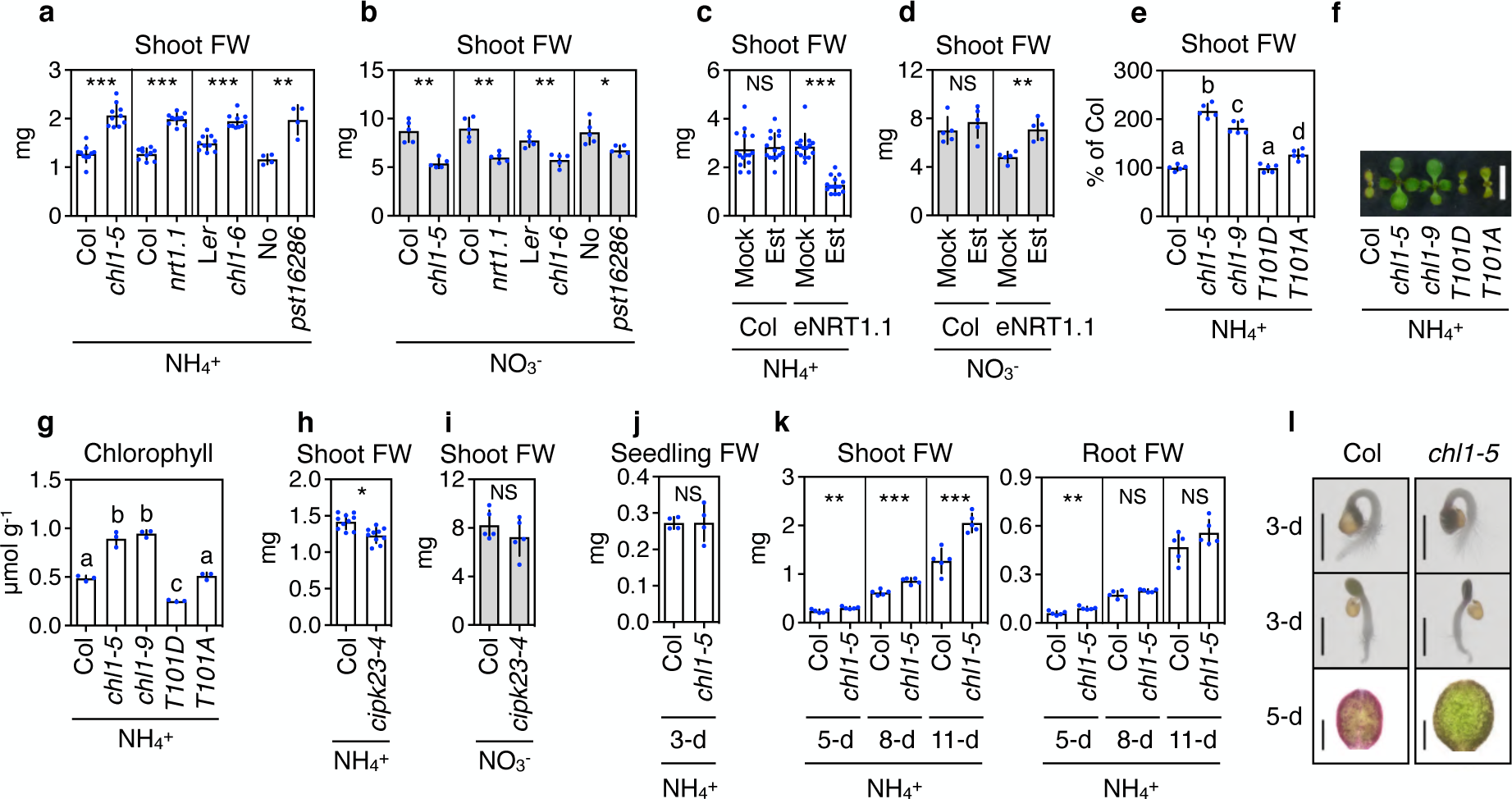
Plasma membrane-targeted NRT1.1 reduces ammonium tolerance almost regardless of its phosphorylating status. (a, b) Fresh weights (FWs) of shoots from 11-day-old Col, *chl1-5*, *nrt1.1*, L*er*, *chl1-6*, Nossen, and *pst16286* plants grown under 10 mM ammonium (a) or 10 mM nitrate (b) conditions (Mean ± SD; n = 5-10). (c, d) Shoot FWs in Col and NRT1.1-mCherry (eNRT1.1) lines under the control of the β-estradiol-inducible promoter in the *chl1-5* background (Mean ± SD; n = 15). In one dish, five seedlings of each line grown under the 10 mM nitrate condition for 5 days were transferred to the 10 mM ammonium (c) or 10 mM nitrate (d) condition in the absence (ethanol as a mock) or presence of 1 µM β-estradiol and further grown for 6 days. (e, f) Shoot FWs (e) and a representative photograph (f) from 11-day-old Col, *chl1-5*, *chl1-9* (*P492L*), *T101D*, and *T101A* plants grown under 10 mM ammonium conditions (Mean ± SD; n = 5). The scale bar represents 5 mm. Twelve shoots from one plate were regarded as a single biological replicate. (g) The chlorophyll (*a* + *b*) concentrations of shoots from 5-day-old Col, *chl1-5*, *chl1-9*, *T101D*, and *T101A* seedlings grown under 10 mM ammonium (Mean ± SD; n = 3). Thirty-seven shoots from one plate were regarded as a single biological replicate. (h, i) FWs of shoots from 11-day-old Col and *cipk23-4* plants grown under 10 mM ammonium (h) or 10 mM nitrate (i) conditions (Mean ± SD; n = 5-10). (j) FWs of 3-day-old Col and *chl1-5* seedlings grown under 10 mM ammonium (Mean ± SD; n = 4). (k) FWs of shoots and roots from 5-, 8-, or 11-day-old Col and *chl1-5* grown under 10 mM ammonium (Mean ± SD; n = 5). (l) Representative photographs of 3-day-old Col and *chl1-5* seedlings grown under 10 mM ammonium and representative photographs of the adaxial side of cotyledons from 5-day-old Col and *chl1-5* seedlings grown under 10 mM ammonium. The scale bars represent 1 mm (3-day) and 500 µm (5-day). (a, b, h–k) Six shoots or roots from one plate were regarded as a single biological replicate. In one dish, six seeds of each line of wild-type and mutant were placed and grown. \**P* < 0.05; \*\**P* < 0.01; \*\*\**P* < 0.001 (unpaired two-tailed Welch’s *t*-test). NS denotes not significant. Different lowercase letters indicate significant differences evaluated by the Tukey–Kramer multiple comparison test conducted at a significance level of *P* < 0.05.

NRT1.1 exhibits diverse molecular functions, including nitrate uptake, nitrate sensing/signaling, and auxin transport (Ho et al. 2009; Krouk et al. 2010; Bouguyon et al. 2015; Zhang et al. 2019). Modifications at specific residues in the protein alter these functions. In the *chl1-9* mutant with the P492L substitution, NRT1.1 primarily localizes intracellularly rather than at the plasma membrane (Bouguyon et al. 2015). Surprisingly, whilst P492L lacks the ability to transport nitrate and auxin, it can still drives some of the nitrate-dependent signaling, including short-term induction of *NRT2.1* by nitrate (Ho et al. 2009; Bouguyon et al. 2015). This suggests that NRT1.1 functions as a nitrate sensor, independent of its uptake activity. The phosphorylation status of NRT1.1 at the T101 residue alters its affinity for nitrate; Non-phosphorylated NRT1.1 forms a homodimer with the NRT1.1 located in close proximity to the dimer interface, allowing low-affinity nitrate uptake, whereas phosphorylated NRT1.1 decouples the dimer configuration to act as a high-affinity nitrate transporter (Sun and Zheng 2015). Moreover, the phosphorylated form transports auxin better than the non-phosphorylated form in the absence of nitrate (Bouguyon et al. 2015). To reveal how P492L substitution and phosphorylation status of NRT1.1 affects ammonium tolerance, we analyzed growth of *chl1-9* and transgenic plants expressing nonphosphomimic (T101A) or phosphomimic (T101D) mutants of NRT1.1.

When grown on 10 mM ammonium, the *chl1-9* mutant exhibited markedly improved ammonium tolerance compared to the wild-type, displaying larger FWs and higher chlorophyll concentrations (Figure 1e–g). Except for slightly enhanced shoot growth in T101A compared to Col, both T101 mutants displayed ammonium toxicity similar to Col. Moreover, *cipk23-4*, lacking CIPK23 that phosphorylates NRT1.1 at the T101 residue (Ho et al. 2009), did not exhibit enhanced shoot growth under ammonium or nitrate conditions (Figure 1h, i). These results imply that NRT1.1-induced ammonium toxicity is nearly independent of its phosphorylation at the T101 residue but requires correct plasma membrane targeting dependent on the P492 residue. Given that the phosphorylating status alters auxin transport in the absence of nitrate (Bouguyon et al. 2015), changes in auxin distribution and action may have little effect on ammonium toxicity.

To determine the developmental stages at which NRT1.1 affects ammonium tolerance, we monitored the growth progression of Col and *chl1-5* plants under 10 mM ammonium. In 3-day-old seedlings, there were no significant difference in FW, and their appearances were similar (Figure 1j, l). By day 5, *chl1*-*5* seedlings exhibited larger shoot and root FWs, and their cotyledons were more expanded and greener compared to Col (Figure 1k, l, and S1a). Additionally, at this stage, *chl1-5* and *nrt1.1* seedlings had significantly higher chlorophyll concentrations than Col (Figure S1b). As plants aged between 8 and 11-days, the impact of *NRT1.1* deficiency on shoot growth intensified compared to root growth (Figure 1k). These findings highlight that NRT1.1’s influence on ammonium tolerance initiates at the early stages of seedling growth.

### 3.2. A significant expression of NRT1.1 is detectable both in shoots and roots of ammonium-grown seedlings

To decipher how NRT1.1 impacts ammonium tolerance, we examined *NRT1.1* expression patterns in Col plants grown under 10 mM ammonium or 10 mM nitrate. In 3-day-old Col seedlings, *NRT1.1* expression showed little disparity between ammonium and nitrate conditions (Figure 2a). However, in 5-, 8-, and 11-day-old plants, shoot expression of *NRT1.1* remained consistently higher under 10 mM ammonium compared to 10 mM nitrate, whereas in the roots, this trend reversed (Figure 2b). Consequently, the disparity in *NRT1.1* transcript levels between shoot and root was smaller under ammonium than nitrate. Furthermore, to investigate NRT1.1 protein expression under ammonium conditions, we utilized the *pNRT1.1:NRT1.1-GFPloop* line for confocal observation (Bouguyon et al. 2016). Clear GFP signals were localized in the plasma membrane of various cells, including pavement cells of cotyledons (Figure 2c, i), epidermal cells of hypocotyl (Figure 2d), mesophyll protoplasts (Figure 2e), root cap cells, and epidermal cells near primary root tips (Figure 2f, g). Intriguingly, dotted GFP signals were frequently detected in intracellular regions of pavement cells and root tips (Figure 2h, j). These expression analyses indicate that NRT1.1 might function in diverse cells and tissues under high ammonium conditions.

**Figure 2.**
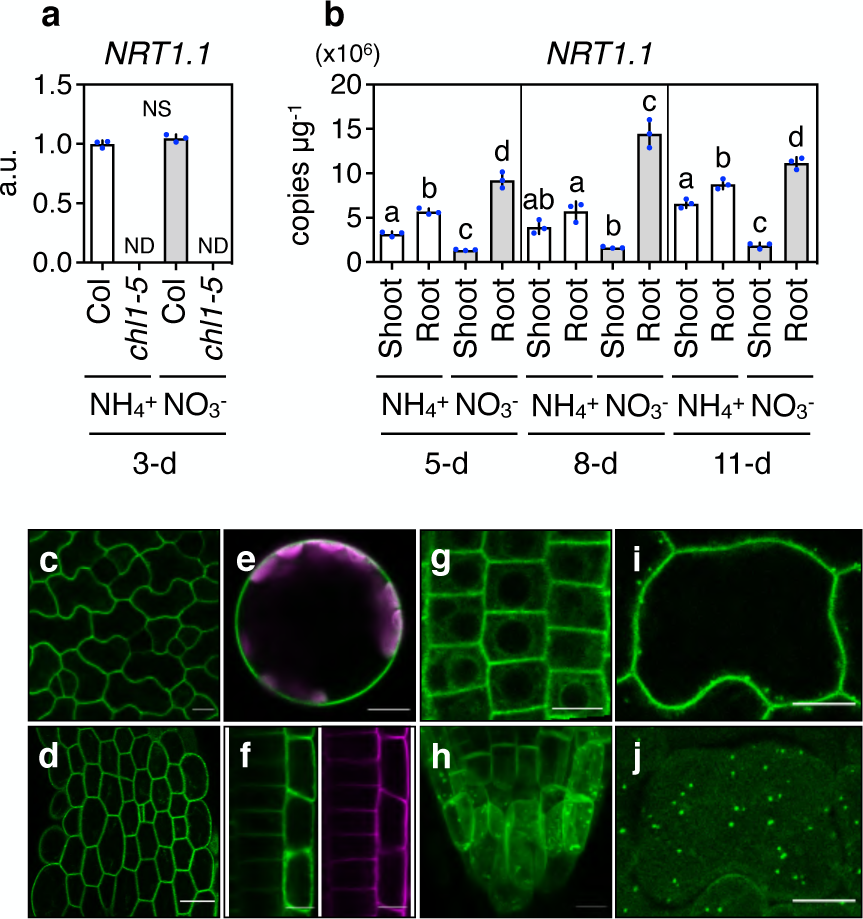
NRT1.1 is expressed in various tissues under ammonium conditions. (a) Relative transcript levels of *NRT1.1* in 3-day-old Col whole seedlings grown under 10 mM ammonium or 10 mM nitrate conditions (Mean ± SD; n = 3). 45 seedlings from one plate were regarded as a single biological replicate. (b) Relative transcript levels of *NRT1.1* in shoots and roots from 5-, 8-, or 11-day-old Col grown under 10 mM ammonium or 10 mM nitrate conditions (Mean ± SD; n = 3). 12 shoots and 12 roots from one plate were regarded as a single biological replicate. (c–j) Signals of NRT1.1-GFPloop from pavement cells of cotyledons from 3-day-old plants (c), epidermal cells of hypocotyl from 3-day-old plants (d), mesophyll protoplasts from 12-day-old plants (e), root cap and epidermal cells near primary root tips from 5-day-old plants (f, g), primary root tip from 5-day-old plants (h), and pavement cells of cotyledons from 8-day-old plants (i: horizontal central section of the cell, j: horizontal section below the cell surface layer) grown under 10 mM ammonium condition except for mesophyll protoplasts from 12-day-old plants (e). Mesophyll protoplasts were prepared from the plants which were grown under 10 mM nitrate for 9 days, then transferred to 10 mM ammonium, and further grown for 3 days. The scale bars represent 10 µm (c, e–j) and 30 µm (d). The purple signals represent autofluorescence (e) and PI fluorescence (f). The images were processed by Image J version 1.53c. Figure 2h was generated by the maximum intensity projection.

### 3.3. Overview of genome-wide transcriptional responses caused by NRT1.1 deficiency

To unravel the primary mechanisms by which NRT1.1 controls genome-wide gene expression under ammonium conditions, we conducted independent microarray experiments using 3-day-old (72 h) and 5-day-old (120 h) whole seedlings of Col, *chl1-5*, and *nrt1.1*. In 3- and 5-day-old seedlings, we identified 20 and 176 transcripts, respectively, in which expression was at least two-fold higher in both *NRT1.1*-deficient mutants compared to Col (Table S2). Conversely, 13 and 80 transcripts in the mutants exhibited levels less than or equal to half of those in Col (Table S2). To capture more NRT1.1-regulated genes in the 3-day-old seedlings, we re-explored differentially expressed genes (DEG) at a 1.5-fold threshold. This analysis identified 120 upregulated genes and 71 downregulated genes in the mutants relative to Col (Table S3). RT-qPCR analysis of 12 DEGs confirmed that the results were consistent with those of the microarray (Figure S2 and Tables S2, 3). For subsequent analyses, we concentrated on DEGs at a 1.5-fold threshold for 3-day-old seedlings and a 2-fold threshold for 5-day-old seedlings. A Venn diagram revealed that only 26 DEGs were common to both 3-day-old and 5-day-old seedlings (Figure 3a and Table S4). Enrichment analysis conducted using Metascape (Zhou et al. 2019) identified a significant enrichment of the term “response to extracellular stimulus” (Figure S3a). Additionally, the SUBA localization predictor (Hooper et al. 2017) indicated that the translational products corresponding to 9 out of the 26 DEGs were likely localized in the extracellular space (Figure S3b). These findings suggest that, under ammonium conditions, NRT1.1 could influence extracellular events.

**Figure 3.**
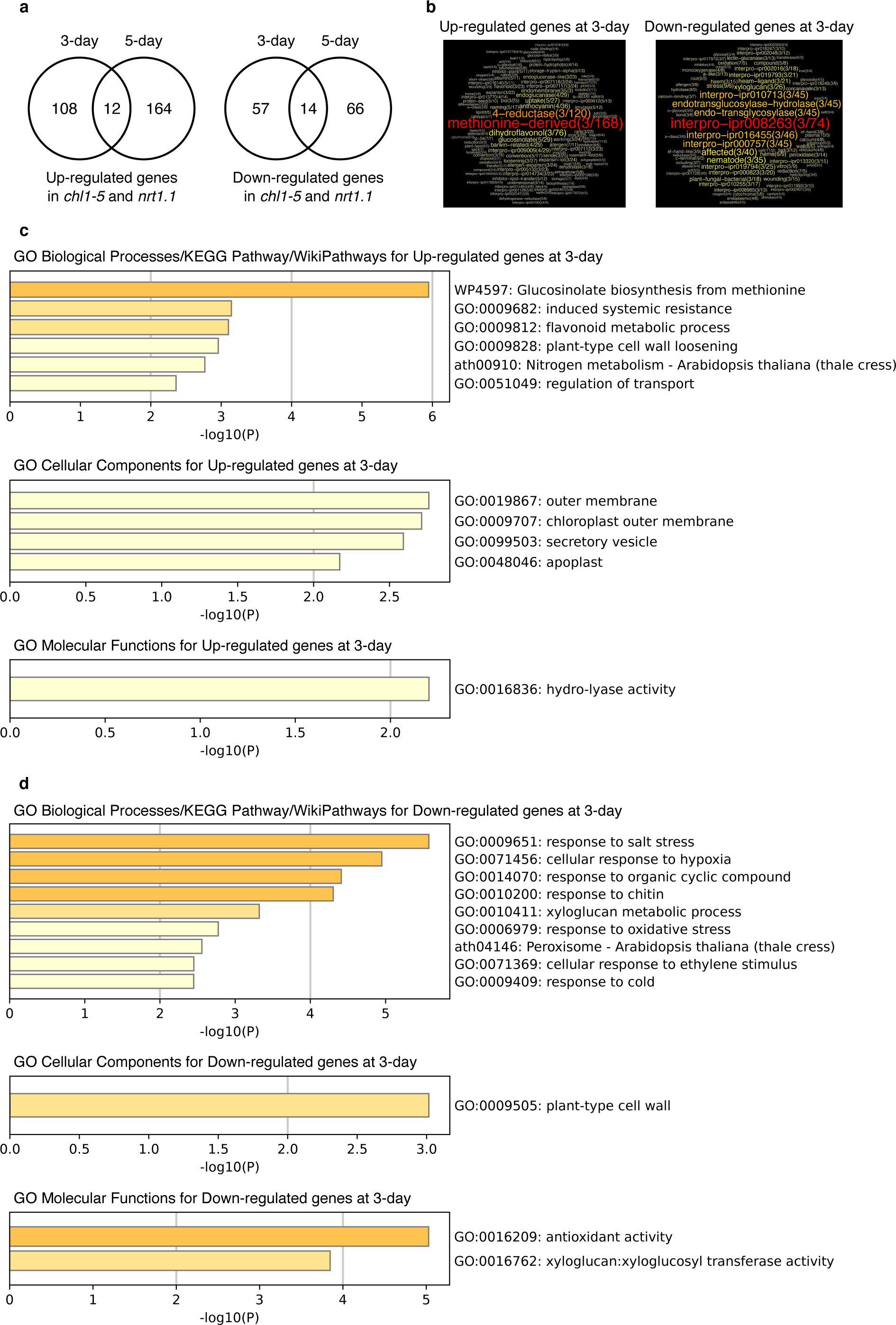
Genome-wide transcriptional responses caused by *NRT1.1* deficiency. (a) Venn diagram showing the number of genes upregulated and downregulated in the 3- and 5-day-old *NRT1.1*-deficient mutants (*chl1-5*, *nrt1.1*) compared with Col grown under 10 mM ammonium. The transcripts whose expression was at least 1.5-fold (3-day) or 2-fold (5-day) higher or at most 1/1.5 (3-day) or 1/2 lower (5-day) in both mutants relative to that in Col were counted. The gene lists are shown in Supplementary Tables S2, 3. (b) Outputs derived from GeneCloud analysis of genes upregulated and downregulated in the 3-day-old *NRT1.1*-deficient mutants (*chl1-5*, *nrt1.1*) compared with Col grown under 10 mM ammonium. The numbers next to the term denote the number of genes containing the term and the fold enrichment. (c, d) Outputs derived from Metascape analysis of genes upregulated (c) and downregulated (d) in the 3-day-old *NRT1.1*-deficient mutants (*chl1-5*, *nrt1.1*) compared with Col grown under 10 mM ammonium. Note that microarray analysis was conducted using 3-day-old (72 h) and 5-day-old (120 h) whole seedlings of Col, *chl1-5*, and *nrt1.1*.

In 3-day-old seedlings, the absence of NRT1.1 led to significant gene expression changes without altering seedling growth (Figure 1e, 3a). Thus, we focused on the transcriptome profile of 3-day-old seedlings to uncover the primary functions of NRT1.1 under ammonium conditions. Gene cloud analysis (Krouk et al. 2015) revealed a 168-fold enrichment in “methionine-derived” and a 120-fold enrichment in “4-reductase” genes in the upregulated genes of 3-day-old *NRT1.1*-deficient mutants (Figure 3b). “Methionine-derived” genes included *At3g19710* (*BCAT4*), *At4g12030* (*BAT5*), and *At5g23010* (*MAM1*), essential for aliphatic glucosinolate biosynthesis. The “4-reductase” genes included *At2g47460* (*MYB12*), *At4g09820* (*bHLH042*), and *At5g42800* (*DFR*), contributing to flavonoid biosynthesis. Metascape analysis also overrepresented terms like “glucosinolate biosynthesis from methionine” and “flavonoid metabolic process” for upregulated genes in the mutants (Figure 3c). Downregulated genes in the 3-day-old mutants enriched terms like “interpro-ipr008263 (glycoside hydrolase, family 16, active site)” from genes encoding cell wall-modifying enzymes like xyloglucan endotransglucosylase/hydrolase (*At4g25810* (*XTH23*), *At4g14130* (*XTH15*), and *At5g57560* (*XTH22*)) (Figure 3b). Intriguingly, DEGs in the 3-day-old mutants often overrepresented terms related to the extracellular space, such as “plant-type cell wall loosening,” “secretory vesicle,” “apoplast,” “xyloglucan metabolic process,” “plant-type cell wall,” and “xyloglucan:xyloglucosyl transferase activity” (Figure 3c, d).

In 5-day-old seedlings, terms related to photosynthesis were predominant among upregulated genes in response to *NRT1.1* deficiency (Figure S4a, b). This aligns with the observed greener and more expanded cotyledons and higher chlorophyll concentrations in *NRT1.1*-deficient seedlings compared to Col (Figure 1m and S1a, b). Conversely, downregulated genes were associated with terms related to abiotic and biotic stresses (Figure S4c). MapMan analysis clearly illustrated the upregulation of genes encoding components for light reactions, chlorophyll biosynthesis, and the Calvin–Benson cycle in 5-day-old *NRT1.1*-deficient mutants (Figure S5b). These findings indicate an enhancement of photoautotrophic growth with reduced stress response in the 5-day-old mutants.

### 3.4. Effects of NRT1.1 deficiency on expression of genes for glucosinolate biosynthesis and genes responsive to ACC, NaCl, H_2_O_2_, IAA, and low pH under ammonium condition

Initially, we concentrated on the genes involved in the biosynthesis of aliphatic GSL and indole GSL, based on the gene list from Harun et al. (2020). This focus was due to the enrichment of terms such as “glucosinolate biosynthesis from methionine,” “methionine-derived,” and “glucosinolate biosynthesis” among the upregulated genes in 3- and 5-day-old *NRT1.1*-deficient mutants (Figures 3b, c, and S5b). The deficiency of *NRT1.1* significantly upregulated the aliphatic GSL biosynthesis genes and their positive regulator genes, *AT5G61420* (*MYB28*) and *AT5G07690* (*MYB29*), while it had little effect on the expression of indole GSL biosynthesis genes (Figure 4a and Tables S2, 3, and 5).

**Figure 4.**
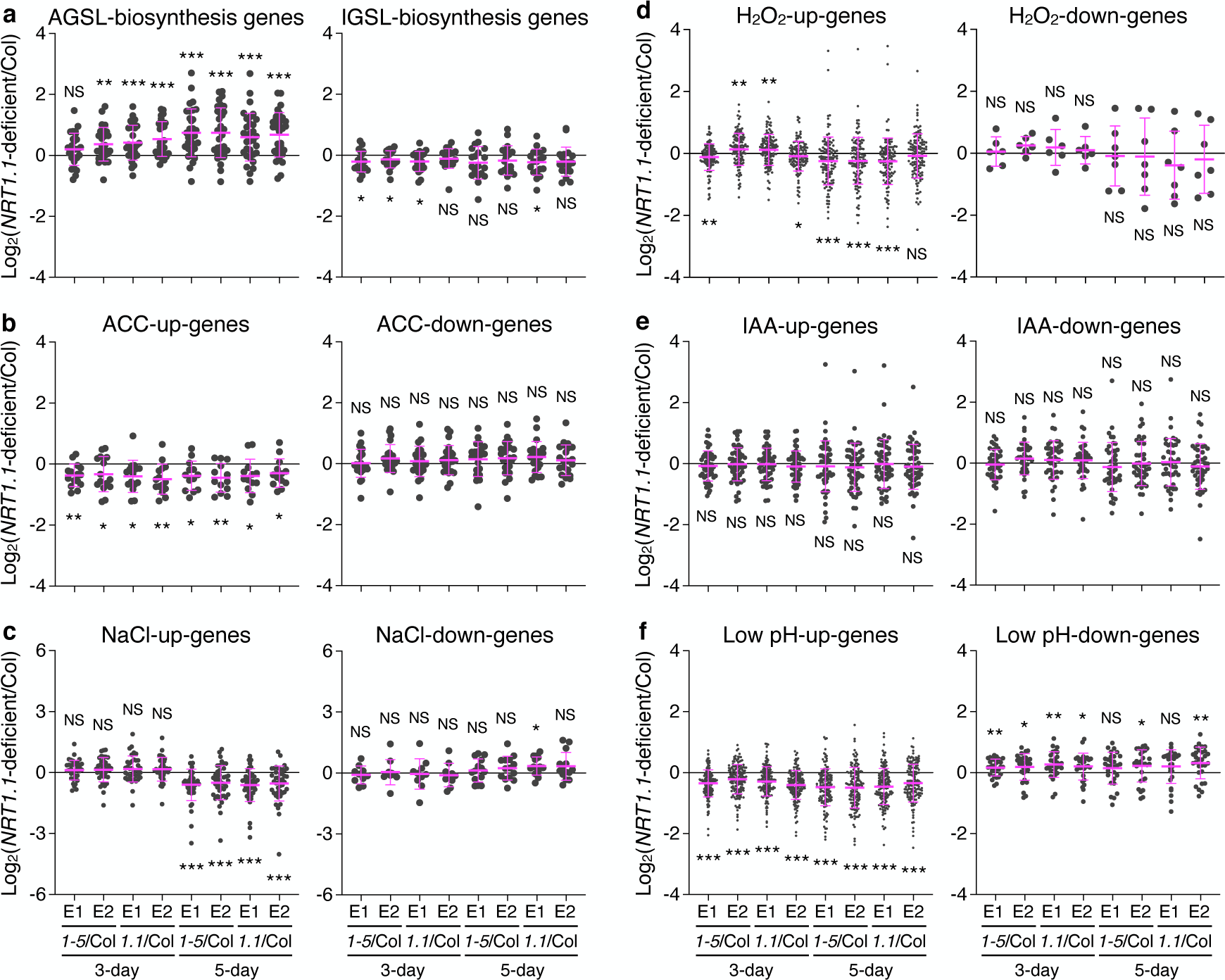
NRT1.1 primarily alters the expression of aliphatic glucosinolate biosynthesis genes, ACC-responsive genes, and low pH-responsive genes under ammonium conditions. (a–f) Comparisons of the expression of biosynthesis genes for aliphatic glucosinolate (AGSL) and indole glucosinolate (IGSL) (a), genes upregulated and downregulated by ACC application (b), genes upregulated and downregulated by NaCl stress (c), genes upregulated and downregulated by H_2_O_2_ application (d), genes upregulated and downregulated by IAA application (e), and genes upregulated and downregulated by low pH treatment (f) between Col and *NRT1.1*-deficient mutants (*chl1-5*, *nrt1.1*) grown under 10 mM ammonium. The samples harvested from two independent experiments were subjected to microarray analysis. E1 and E2 denote 1st experiment and 2nd experiment, respectively. One hundred thirty-five seedlings (3-day) from three plates and 74 seedlings (5-day) from two plates were regarded as a single biological replicate. Changes in gene expression levels between Col and *chl1-5* and Col and *nrt1.1* were represented as logarithms to base 2 of ratios in signal intensities. *1-5* and *1.1* denote *chl1-5* and *nrt1.1*, respectively. \**P* < 0.05; \*\**P* < 0.01; \*\*\**P* < 0.001 (two-tailed ratio paired *t*-test).

Our enrichment analysis revealed that the GO term “cellular response to ethylene stimulus” was overrepresented among the downregulated genes in 3-day-old *NRT1.1*-deficient mutants (Figure 3d). This aligns with a previous study that reported a down-regulation of genes involved in ethylene biosynthesis and senescence under toxic ammonium conditions in *NRT1.1*-deficient mutants (Jian et al. 2018). Consequently, we examined the genes responsive to the application of 1-aminocyclopropane-1-carboxylic acid (ACC), a precursor of ethylene whose biosynthesis is a rate-limiting step for ethylene production, based on the gene list from Goda et al. (2008) (Figure 4b and Table S6). The ACC-induced genes were significantly downregulated in both 3- and 5-day-old *NRT1.1*-deficient mutants grown under 10 mM ammonium, whereas the ACC-repressed genes did not show a significant response.

The GO term “response to salt stress” was found to be overrepresented in the downregulated genes of 3- and 5-day-old *NRT1.1*-deficient mutants (Figures 3d and S4c). A recent study found that in the presence of a 25 mM chloride ion with ammonium as the sole N source, NRT1.1 supports excessive chloride absorption, resulting in severe root growth inhibition (Liu et al. 2020). *NRT1.1* deficiency may affect the expression of NaCl-responsive genes because our ammonium solution contains 16 mM chloride ions. The NaCl-induced genes listed from Shen et al. (2014) were significantly downregulated in the 5-day-old *NRT1.1*-deficient mutants, but not in the 3-day-old mutants (Figures 4c and Table S7). The expression of NaCl-repressed genes differed little between Col and *NRT1.1*-deficient mutants.

It has been reported that ammonium nutrition disrupts redox homeostasis, leading to the apoplastic accumulation of reactive oxygen species and oxidative stress (Podgórska et al. 2015). In line with this, our enrichment analysis revealed the presence of the GO terms “response to oxidative stress” and “antioxidant activity” among the downregulated genes in both 3-day-old and 5-day-old *NRT1.1*-deficient mutants (Figure 3d and S4c). Genes that are upregulated in response to H_2_O_2_ application, as documented by Hieno et al. (2019), were downregulated in the 5-day-old *NRT1.1*-deficient mutants, but not in the 3-day-old mutants (Figure 4d and Table S8). No significant alterations in the expression of genes downregulated by H_2_O_2_ were observed between Col and the *NRT1.1*-deficient mutants.

Regarding auxin responses, although NRT1.1 facilitates auxin influx into the cell in the absence of nitrate (Krouk et al. 2010), its deficiency did not significantly alter the expression of typical genes responsive to indole-3-acetic acid (IAA) application (Goda et al. 2008) (Figure 4e and Table S9).

Interestingly, previous studies have shown that 43% of ammonium-inducible genes correspond to low pH (4.5)-inducible genes in Arabidopsis plants (Lager et al. 2010; Patterson et al. 2010). We have previously observed that the enhanced ammonium tolerance of *NRT1.1*-deficient mutants at pH 5.7 was mimicked in Col plants grown at pH 6.7 (Hachiya et al. 2011). These findings suggest that *NRT1.1*-deficient mutants may exhibit enhanced tolerance to ammonium-derived acidic stress. Consequently, we examined the microarray data with a focus on low pH stress-responsive genes listed in Lager et al. (2010). Remarkably, the low pH stress-inducible genes were significantly downregulated both in the 3- and 5-day-old *NRT1.1*-deficient mutants relative to Col, whereas the low pH stress-repressive genes were generally upregulated in the mutants (Figure 4f and Table S10).

Collectively, we conclude that NRT1.1 primarily alters the expression of genes associated with aliphatic glucosinolate biosynthesis, ethylene signaling, and low pH stress.

## 4. Discussion

The Arabidopsis nitrate transceptor NRT1.1 and its orthologs are required for plant adaptation to nitrate-rich conditions (Wang et al. 2020). NRT1.1, on the other hand, has been shown to reduce ammonium tolerance in the absence of nitrate (Hachiya et al. 2011; Jian et al. 2018). Thus, NRT1.1 and its orthologs are likely to determine adaption features for the two major N sources, nitrate and ammonium.

Excessive ammonium assimilation by plastidic glutamine synthetase produces proton accumulation and acidic stress in Arabidopsis shoots when exposed to high amounts of ammonium (Hachiya et al. 2021). Proton excretion from plants to external media is common in plant cultivation with high ammonium feeding (Britto and Kronzucker 2002). Moreover, in Arabidopsis plants, ammonium-inducible genes and low pH (4.5)-inducible genes overlap at a significant rate (Lager et al. 2010, Patterson et al. 2010). These findings imply that acidic stress is a major source of ammonium toxicity. Our transcriptome analysis revealed that *NRT1.1* deficiency dampens low pH responses (Figure 4f). We observed that increasing the pH of the medium from 5.7 to 6.7 by adding alkaline NH_3_ solution promoted shoot development of Col more intensely than that of *chl1-5* under ammonium conditions, virtually completely suppressing the difference in growth (Figure S6a). Expression of *ALMT1* was upregulated in Col shoots and roots under 10 mM ammonium compared with under 10 mM nitrate, and this ammonium induction was significantly suppressed in *chl1-5* shoots and roots (Figure S6b). Importantly, the magnitude of ammonium induction and suppression by *NRT1.1* deficiency of *ALMT1* was much larger in shoots than in roots (Figure S6b). This corresponds to the observation that the impact of *NRT1.1* deficiency on shoot growth intensified compared to root growth (Figure 1k). These suggest that NRT1.1 exacerbates acidic stress in the presence of ammonium, mainly in shoots. Interestingly, an acidic rhizosphere induces *NRT1.1* expression in a STOP1 transcription factor-dependent way, which enhances symport of proton and nitrate through NRT1.1 and adjusts rhizosphere pH to a more suitable range (Ye et al. 2021). Furthermore, NRT1.1 and SLAH3 collaborate to drive a nitrate cycle across the plasma membrane, reducing acidification of the rhizosphere (Xiao et al. 2022). It is uncertain how NRT1.1 affects intra- and extracellular pH under nitrate-free ammonium conditions.

Our transcriptome analysis highlighted the extracellular space as an early site of NRT1.1 action (Figure 3c, d, and S3a, b, S4b, c). A study by Podgórska et al. (2017) in Arabidopsis plants demonstrated that ammonium toxicity is associated with smaller mesophyll cells characterized by a more rigid cell wall structure, containing increased phenolic compounds and boron ions. In our study, the GO terms “plant-type cell wall loosening” and “plant-type cell wall modification” were enriched in the genes upregulated by *NRT1.1* deficiency (Figure 3c and S4b). These terms originated from genes encoding proteins involved in cell wall loosening and cell expansion, such as *AT1G69530* (*EXPA1*), *AT1G74670* (*GASA6*), *AT2G20750* (*EXPB1*), *AT2G40610* (*EXPA8*), *AT3G29030* (*EXPA5*), and *AT4G28250* (*EXPB3*). This suggests that NRT1.1 might influence ammonium tolerance via alterations in cell wall modification processes. We discovered that genes for aliphatic GSL biosynthesis and their positive regulator genes, *MYB28* and *MYB29*, were induced by *NRT1.1* deficiency (Figure 4a, S2b, Table S2, 4). Coleto et al. (2021) reported that ammonium toxicity was exacerbated in the *myb28myb29* mutant but not in the *myc234* mutant. Given that both mutants are almost devoid of aliphatic GSL, the ammonium hypersensitivity of *myb28myb29* is not linked with a lack of aliphatic GSL. The authors provided evidence that MYB28 and MYB29 maintain intracellular iron homeostasis under ammonium conditions, thereby attenuating ammonium toxicity. It remains to be seen whether the absence of NRT1.1 enhances ammonium tolerance via upregulation of *MYB28* and *MYB29*. Meanwhile, our transcriptome data suggested that NRT1.1 enhances ethylene signaling under ammonium conditions. Plants emit ethylene as a phytohormone in response to various stresses, including ammonium toxicity (Britto and Kronzucker 2002). Inhibition of ethylene signaling by gene knockout and chemical treatment alleviates ammonium toxicity (Jian et al. 2018, Li et al. 2019). These findings suggest that NRT1.1 reduces ammonium tolerance through ethylene signaling. Although there is little overlap among aliphatic GSL biosynthesis genes and ACC- and low pH-responsive genes (Tables S5, 6, 10), it would be worthwhile to scrutinize these associations.

## Acknowledgments

We are grateful for Dr. Carine Alcon and Montpellier RIO Imaging on technical support with confocal microscopy and for Ms. Mako Sakai on technical support with expression analysis and on manuscript review. We would like to thank Dr. Yi-Fang Tsay (Academia Sinica) for giving the Arabidopsis seeds. We would also like to thank Enago (www.enago.jp) for the English language review.

