## Supplementary Figures for "Genetic and transcriptomic dissection of nitrate-independent function of Arabidopsis NRT1.1/NPF6.3/CHL1 under high ammonium condition"

### Supplementary Figure S1

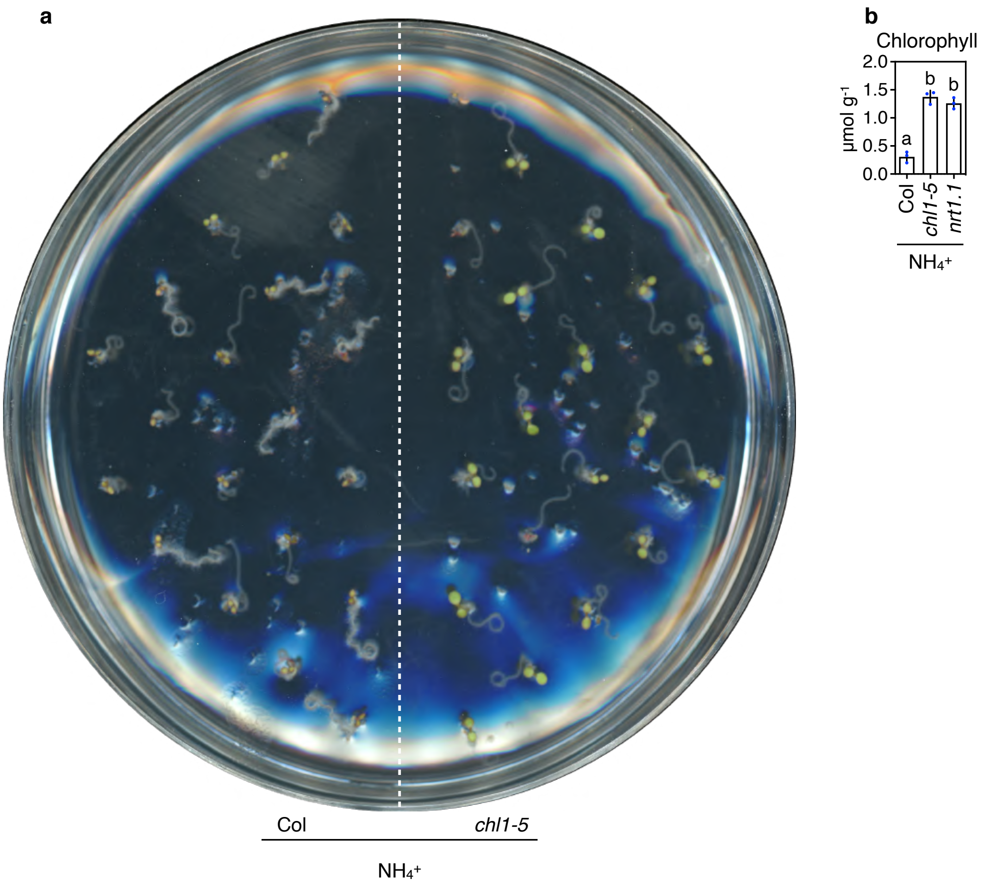


**Fig. S1** Phenotypes of 5-day-old Col and *NRT1.1*-deficient mutants grown under 10 mM ammonium. (a) A representative photograph from 5-day-old Col and *chl1-5*. (b) The chlorophyll (*a* + *b*) concentrations of shoots from 5-day-old Col, *chl1-5*, and *nrt1.1* grown under 10 mM ammonium (Mean ± SD; n = 3). 37 shoots from one plate were regarded as single biological replicate.

### Supplementary Figure S2

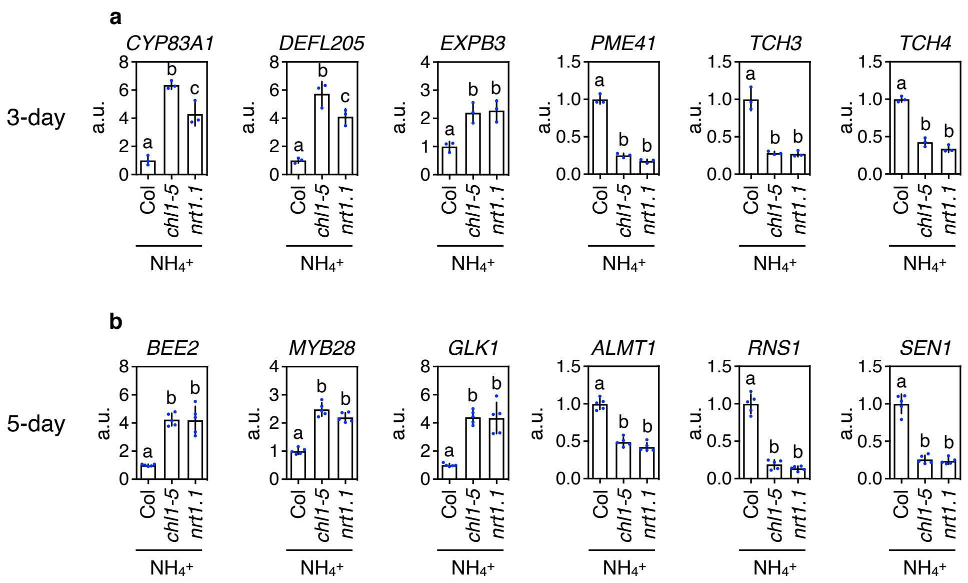


**Fig. S2** Transcriptional responses caused by *NRT1.1* deficiency. (a) Relative transcript levels of *CYP83A*, *DEFL205*, *EXPB3*, *PME41*, *TCH3*, and *TCH4* in 3-day-old Col and *NRT1.1*-deficient mutants (*chl1-5*, *nrt1.1*) grown under 10 mM ammonium (Mean ± SD; n = 3). Forty-five seedlings from one plate were regarded as single biological replicate. (b) Relative transcript levels of *BEE2*, *MYB28 GLK1*,*ALMT1*, *RNS1*, and *SEN1* in 5-day-old Col, *chl1-5*, and *nrt1.1* grown under 10 mM ammonium (Mean ± SD; n = 5). Thirty-seven seedlings from one plate were regarded as single biological replicate.

### Supplementary Figure S3

**a**

GO Biological Processes/KEGG Pathway/WikiPathways for DEGs common to 3-day- and 5-day-old seedlings

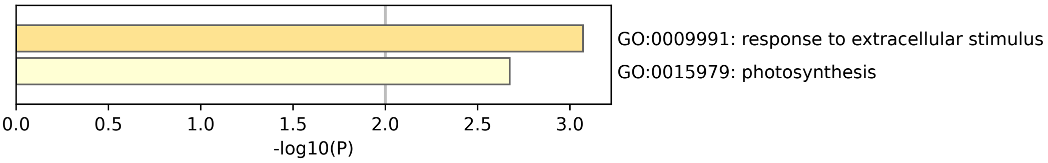


**b**

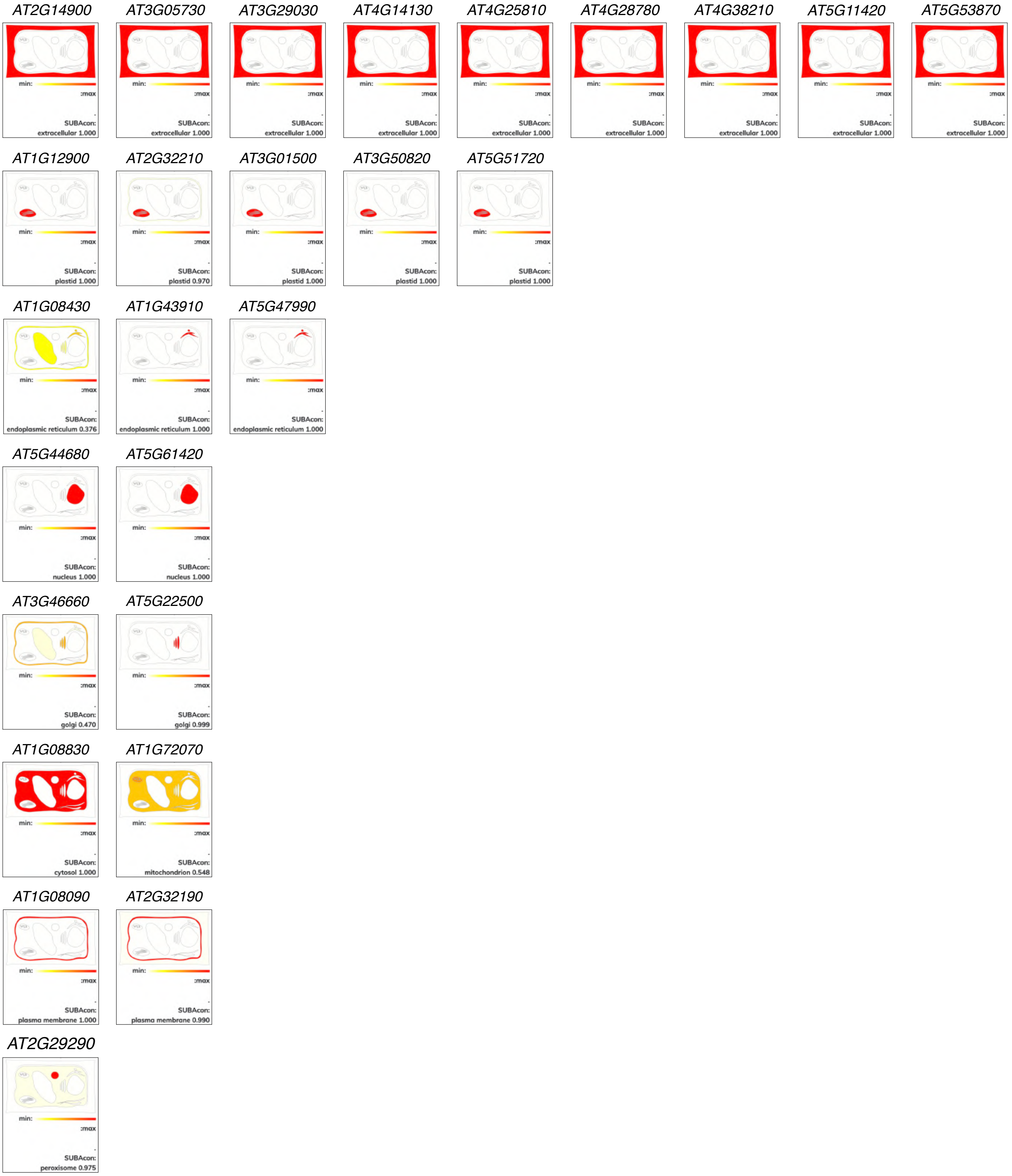


**Fig. S3** Analyses of 25 DEGs common to the 3-day-old and 5-day-old seedlings (Figure 3a and Table S4). (a) Outputs derived from Metascape (Zhou et al. 2019) analysis of the DEGs. (b) Subcellular localization of the DEGs-encoded proteins predicted by SUBA5 (https://suba.live).

### Supplementary Figure S4

**a**

Up-regulated genes at 5-day    Down-regulated genes at 5-day

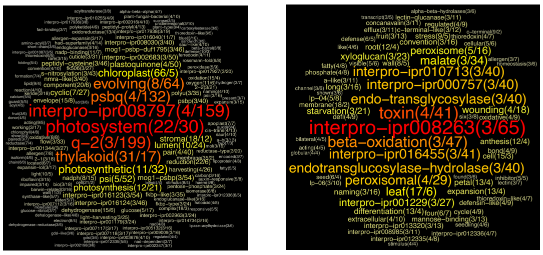


**b**

GO Biological Processes/KEGG Pathway/WikiPathways for Up-regulated genes at 5-day

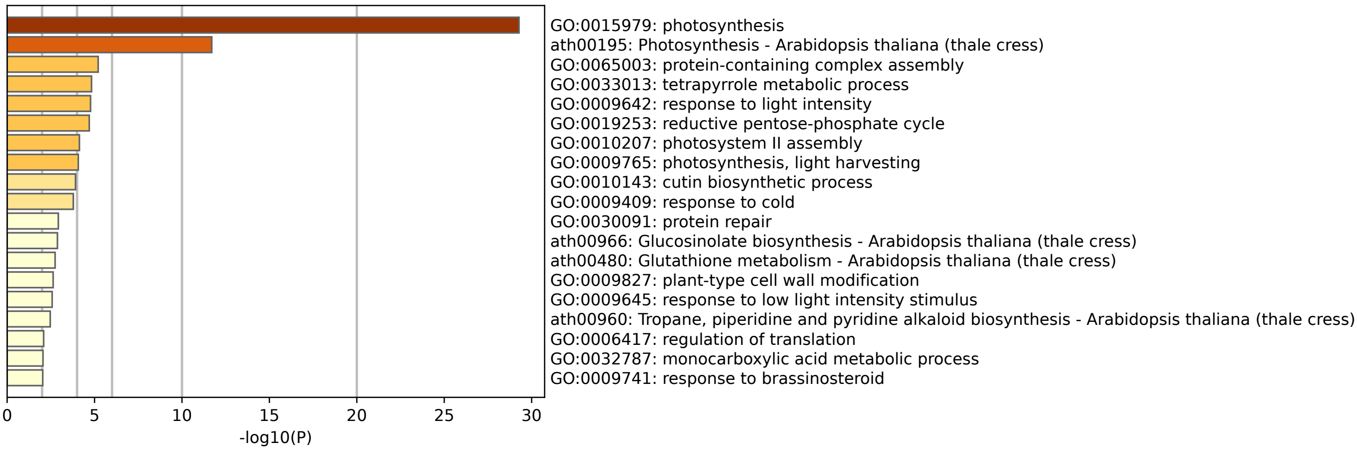


**c**

GO Biological Processes /KEGG Pathway /WikiPathways for Down-regulated genes at 5-day

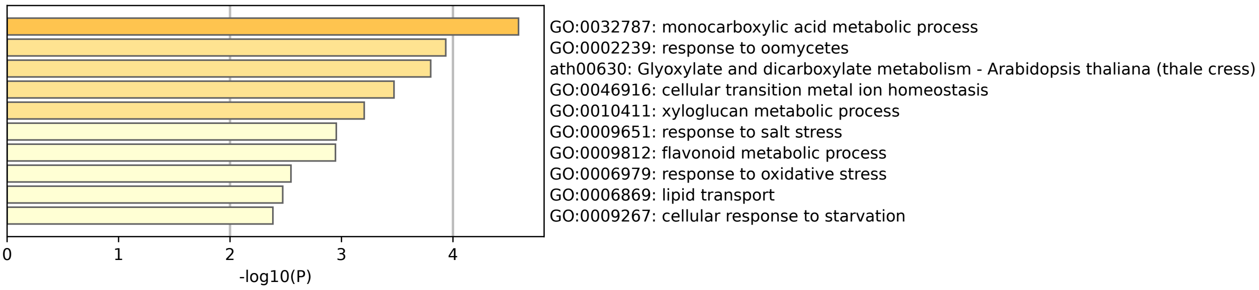


**Fig. S4** Genome-wide transcriptional responses caused by *NRT1.1* deficiency. (a) Outputs derived from GeneCloud (Krouk et al. 2015) analysis of genes upregulated or downregulated in the 5-day-old *NRT1.1*-deficient mutants (*chl1-5*, *nrt1.1*) compared with Col grown under 10 mM ammonium. The numbers next to the term denote the number of genes containing the term and the fold enrichment. (b, c) Outputs derived from Metascape (Zhou et al. 2019) analysis of genes upregulated (b) or downregulated (c) in the 5-day-old *NRT1.1*-deficient mutants (*chl1-5*, *nrt1.1*) compared with Col grown under 10 mM ammonium.

**Supplementary Figure S5**

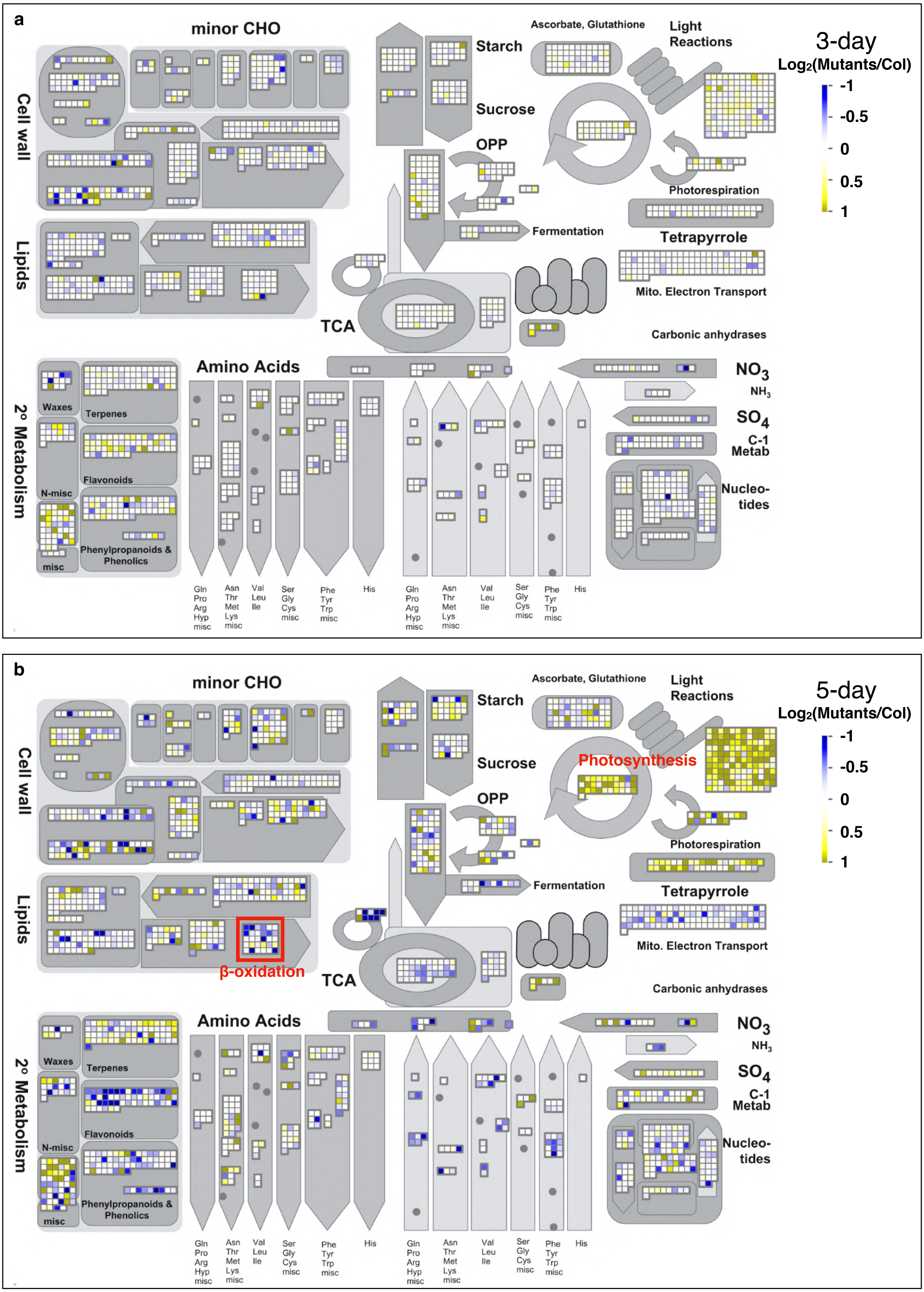


**Fig. S5** Outputs from MapMan analysis of gene expression in the (a) 3- and (b) 5-day-old *NRT1.1*-deficient mutants (*chl1-5*, *nrt1.1*) compared with Col grown under 10 mM ammonium condition. For the comparison between Col and the mutants, the mean values of the transcript levels in *chl1-5* and *nrt1.1* were used, and the values of 1st and 2nd experiments were also averaged.

### Supplementary Figure S6

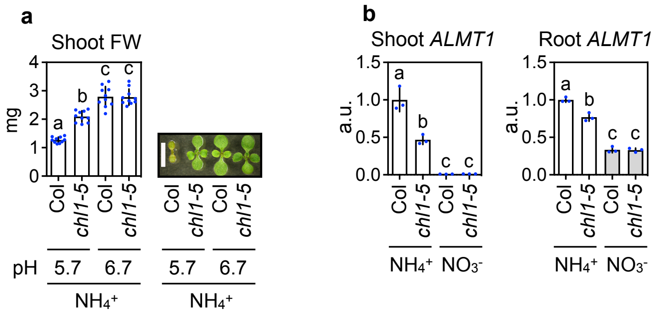


**Fig. S6** NRT1.1 exacerbates acidic stress under ammonium conditions. (a) Effects of medium pH on shoot FWs of 11-day-old Col and *chl1-5* grown under 10 mM ammonium conditions (Mean ± SD; n = 10). Six shoots from one plate were regarded as a single biological replicate. The pH was adjusted to 5.7 by 1N KOH solution, and then alkaline ammonia solution for further pH adjustment from 5.7 to 6.7. The scale bar represents 5 mm. (b) Relative transcript levels of *ALMT1* in shoots and roots from 5-day-old Col, *chl1-5*, and *nrt1.1* grown under 10 mM ammonium (Mean ± SD; n = 5). Thirty-seven seedlings from one plate were regarded as single biological replicate.
